# Prediction and scanning of IL-5 inducing peptides using alignment-free and alignment-based method

**DOI:** 10.1101/2022.10.19.512965

**Authors:** Naorem Leimarembi Devi, Neelam Sharma, Gajendra P. S. Raghava

## Abstract

Interleukin-5 (IL-5) is the key cytokine produced by T-helper, eosinophils, mast and basophils cells. It can act as an enticing therapeutic target due to its pivotal role in several eosinophil-mediated diseases. Though numerous methods have been developed to predict HLA binders and cytokines-inducing peptides, no method was developed for predicting IL-5 inducing peptides. All models in this study have been trained, tested and validated on experimentally validated 1907 IL-5 inducing and 7759 non-IL-5 inducing peptides obtained from IEDB. First, alignment-based methods have been developed using similarity and motif search. These alignment-based methods provide high precision but poor coverage. In order to overcome this limitation, we developed machine learning-based models for predicting IL-5 inducing peptides using a wide range of peptide features. Our random-forest model developed using selected 250 dipeptides achieved the highest performance among alignment-free methods with AUC 0.75 and MCC 0.29 on validation dataset. In order to improve the performance, we developed an ensemble or hybrid method that combined alignment-based and alignment-free methods. Our hybrid method achieved AUC 0.94 with MCC 0.60 on validation/ independent dataset. The best model developed in this study has been incorporated in the web server IL5pred (https://webs.iiitd.edu.in/raghava/il5pred/).

**Key Points:**

- IL-5 is a regulatory cytokine that plays a vital role in eosinophil-mediated diseases
- BLAST-based similarity search against IL-5 inducing peptides was employed
- A hybrid approach combines alignment-based and alignment-free methods
- Alignment-free models are based on machine learning techniques
- A web server ‘IL5pred’ and its standalone software have been developed

**Authors’ Biography:**

1. Dr. Naorem Leimarembi Devi is currently working as a DBT-Research Associate in Department of Computational Biology, Indraprastha Institute of Information Technology, New Delhi, India.
2. Neelam Sharma is pursuing her Ph.D. in Computational Biology from the Department of Computational Biology, Indraprastha Institute of Information Technology, New Delhi, India.
3. Prof. G.P.S. Raghava is currently working as Professor and Head of Department of Computational Biology, Indraprastha Institute of Information Technology, New Delhi, India.

## Introduction

Innate immunity is considered the first line of defense against invading pathogens [1]. In contrast, adaptive immunity act as the second line of defense and is referred to as antigen-specific immune response. The immunological response against a pathogen is mediated via major histocompatibility complex (MHC) by presenting the antigen on the surface of antigen-presenting cells including dendritic, B-cells and macrophages [1]. MHC class II is responsible for processing and presenting exogenous antigens. This antigenic peptide activates CD4+ T-helper cells (Th) hence inducing the release of different cytokines [2]. Interleukin-5 (IL-5) is a crucial cytokine secreted by Th2, type-2 innate lymphoid, mast, basophils and eosinophils cells [3,4]. These cells produced IL-5 upon stimulation by several environmental pollutants, inhaled allergens and microbes [5]. IL-5 is a glycosylated homodimeric protein with 45-60 kDa molecular weight, composed of two helical bundle motifs [6]. The gene that encodes IL-5 is found in the same cluster as IL-3, IL-4, IL-13 and granulocyte-macrophage colony-stimulating factor (GM-CSF) [7]. IL-5 exerts its pleiotropic actions via IL-5 receptor (IL-5R), which is comprised of an α and a βc chain. The α subunit recognises the IL-5 molecule, whereas the βc subunit recognises either IL-3 or GM-CSF. Thereby promoting eosinophils maturation, activation, survival and discharge from the bone marrow into the bloodstream and finally to the airways [5,8]. When activated by IL-5, eosinophils degranulate and produce antimicrobial cytotoxins that are harmful to nearby cells and tissues [9]. Although IL-5 is critical in eosinophil development, it has also been linked to the onset and severity of a number of diseases, including asthma, autoimmune disorders, allergy, atopic dermatitis, eosinophilic esophagitis, and cancer [5,10–13]. IL-5 production during asthma exacerbation can cause pulmonary eosinophilia, enhancing airway smooth muscle contraction and increased mucus production [14]. It has been demonstrated in earlier studies that IL-5 is induced by a wide variety of MHC class II alleles and has a role in MHC class II regulation [15–17]. The overall biological effects of IL-5 on eosinophils are depicted in Figure 1.

**Figure 1:**
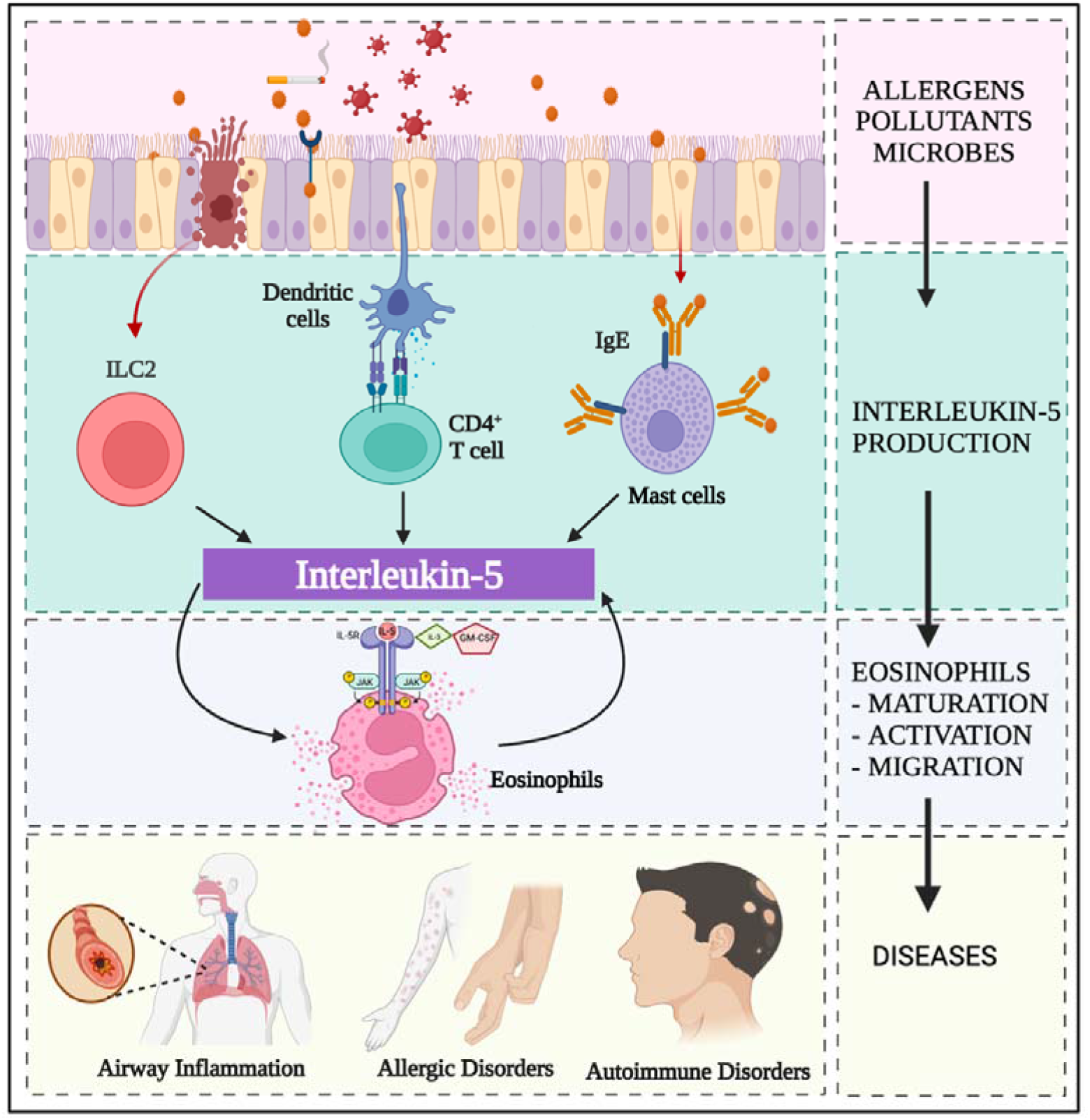
The biological effects of IL-5 on eosinophils

In the past, numerous computational methods have been developed for designing peptide/epitope-based subunit vaccines and immunotherapy. In the initial phase, methods have been developed for predicting Human Leukocyte Antigen (HLA) class I and II binders. Major HLA class I binder prediction include ProPred1 [18], nHLAPred [19], PSSMHCpan [20], NetMHCpan [21] and NetMHC-3.0 [22]. Major HLA class II binder prediction methods include ProPred [23], HLA-DR4Pred [24], NetMHCII [25] and NetMHCIIpan [26]. Besides, another method HLA_nc_Pred [27] has been developed to predict non-classical HLA binders. Recently, the trend has changed as a number of researchers are developing methods for predicting peptides that can induce specific types of cytokines. CytoPred is a method that can predict and classify cytokines with high accuracy [28]. Currently, the following methods, IL4pred [29], IL-6Pred [30], IL-10Pred [31], IL17eScan [32], and IL-13Pred [33] have been developed to predict IL-4, IL-6, IL-10, IL-17, and IL-13 inducing peptides respectively. Moreover, PIP-EL [34] and ProInflam [35] are the tools used to predict the peptides inducing a group of cytokines.

To the best of our knowledge, no method has been developed to predict IL-5 inducing peptides. Thus in this study, we proposed a novel method IL5pred to identify these peptides. We retrieved the experimentally validated dataset from the well-established database Immune Epitope Database (IEDB) [36]. We have implemented Basic Local Alignment Search Tool (BLAST) to predict the peptides based on their similarity to known IL-5 inducers. A motif-based approach was utilised to identify motifs in IL-5 inducers. Next, a wide range of sequence-based features for each peptide was computed using Pfeature tool [37]. Finally, several machine learning models were developed to predict IL-5 inducers and the best model was incorporated into the web server that can efficiently classify IL-5 inducers and non-inducers.

## Materials and Methods

### Dataset preparation

The most challenging aspect of designing a bioinformatics tool is gathering enough experimentally validated data. The current study retreived IL-5 inducing and non-IL-5 inducing peptides from IEDB [36], a freely accessible database that consists of a huge amount of experimentally verified immune epitopes. We created two datasets: main dataset and alternate dataset.

#### Main dataset

We extracted 2802 IL-5 inducing peptides and 8674 non-IL-5 inducing peptides, which are experimentally validated MHC class II binders from IEDB tested on *Homo sapiens*. Further, all peptides containing non-standard characters (i.e., ‘B,’ ‘J,’ ‘O’, ‘U,’ ‘X’ and ‘Z’) and duplicates were removed. We have chosen unique linear peptides with lengths ranging from 9-20 amino acids. The peptides found to be common or exactly matched in both datasets were also eliminated. Finally, 1907 IL-5 inducing and 7759 non-IL-5 inducing peptides were obtained and labelled as positive and negative datasets, respectively.

#### Alternate dataset

We have also created an alternate dataset comprising IL-5 inducing as a positive dataset and random peptides as a negative dataset (generated from the Swiss-Prot database) [38]. We randomly extracted 1907 peptides and assigned them as negative peptides to create a balanced dataset. Finally, the alternate dataset comprises 1907 IL-5 inducing peptides and 1907 random peptides.

### Sequence logos

To investigate the preference of individual amino acids at a particular position, we generated the sequence logo using the R package “ggseqlogo” [39]. It gives the graphical representation with residue positions on x-axis and the bit score indicating the conservation of residues at a specific position on y-axis. This package processes a fixed-length input vector, thus, we considered the minimum length of 9 amino acids from N-terminal for each peptide sequence.

### Motif identification using MERCI

The detection of motifs in peptides is crucial for annotating the function of the sequence. The current study employed a publicly available software, Motif-EmeRging and with Classes-Identification (MERCI) [40] to search motifs in IL-5 inducing peptides. This software used a Perl script to find the motifs exclusively present in positive and negative peptide sequences.

### Feature generation

Pfeature [37], a standalone tool, was used to generate distinct features from peptide sequence information. Using this tool, we computed several compositional-based features as well as binary profile-based features for each peptide sequence. The major composition features such as amino acid composition (AAC), dipeptide composition (DPC), tripeptide composition (TPC), atom composition (ATC), Physico-chemical properties repeat composition (PRI) along with their vector length are tabulated in Table S1. Following are some of the descriptions of the composition-based features used in the current study to develop several alignment-free prediction models using machine learning (ML) techniques.

### Amino acid composition-based features

AAC has been used efficaciously in a number of sequence-based classification techniques [41]. It is the primary feature that describes the fraction of each amino acid residue present in a peptide sequence with 20 length vector. Equation (1) is used to calculate AAC for each amino acid residue.

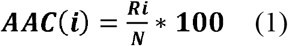

AAC(i) represents the per cent amino acid composition of residue type (i); Ri represents the number of residues of type i, and N represents the length of the sequence.

### Dipeptide composition-based features

DPC is another frequently utilised input parameter for peptide composition-based classification. It captures comprehensive information about the pairwise composition of the amino acids in the peptide sequence with a fixed vector of length 400 (20*20). The following equation is used to calculate DPC.

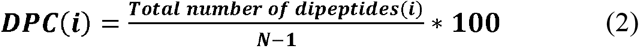

DPC(i) represents the per cent of dipeptide composition of residue type i, and N represents the length of the sequence.

### Selection and ranking of features

A total of 9553 features were computed for both main and alternate datasets. Only relevant features are used for developing the classification model. Thus, selecting relevant features from a larger set of features is the most important step but challenging. Although there are a number of techniques for feature selection, we employed a support vector classifier (SVC)-L1 (Scikit-learn package) to choose the small set of relevant features [42]. Feature-selector tool was further used to rank these features based on their importance. This tool rank features that are frequently used to divide the dataset among all trees, using the DT-based algorithm Light Gradient Boosting Machine [43].

### Binary profile-based features

Previous research studies have shown that the binary profile is one of the essential features for classifying peptides [44]. Generally, it is challenging to create a fixed-length pattern given the varied length of the peptides. In our study, the length of the peptides varies from 9-20, thus we generated a fixed length binary profile by extracting the fixed length segments from either N or C terminus of the peptide [44]. Since the minimum length of the peptides is 9, thus we created binary profiles for N_9_, C_9_ and also for combined terminal residues N_9_C_9_ after calculating the fixed length patterns.

### Alignment-based search using BLAST

In this study, BLAST has been used for alignment-based search. It is the most frequently used tool for annotating protein/peptide and nucleotide sequences [45]. We implemented blastp-short for short peptide sequences (8-30 amino acids) to identify IL-5 inducing peptides based on the similarity of peptides with IL-5 inducers and non-inducers. To identify IL-5 inducers and non-inducers, the top hit of BLAST was considered at different E-value cut-offs.

### Alignment-free approach for classification

To generate more accurate prediction methods, several studies have employed alignment-free approach using different ML techniques [46,47]. For this, Random Forest (RF) [48], K-nearest neighbour (KNN) [49], Decision Tree (DT) [50], Gaussian Naïve Bayes (GNB) [51], XGBoost (XGB) [52], and Logistic Regression (LR) [53] were employed using Scikit’s sklearn package from Python [42]. Different hyperparameters were tuned in these classification algorithms, and the results achieved on the best parameters were reported.

### Five-fold cross-validation

The datasets were split into 80:20 ratio, in which 80% of the data was utilised for training and 20% for validation purposes. Five-fold cross-validation (CV) was used on 80% training dataset to train, test and evaluate the classification models. The training dataset is split into five equal folds, four of which are utilised for training and the remaining for testing. This method is iterated five times, with each fold being tested separately. This method has been extensively applied in several studies by researchers in the past [44,54]. The complete workflow of IL5pred is shown in Figure 2.

**Figure 2:**
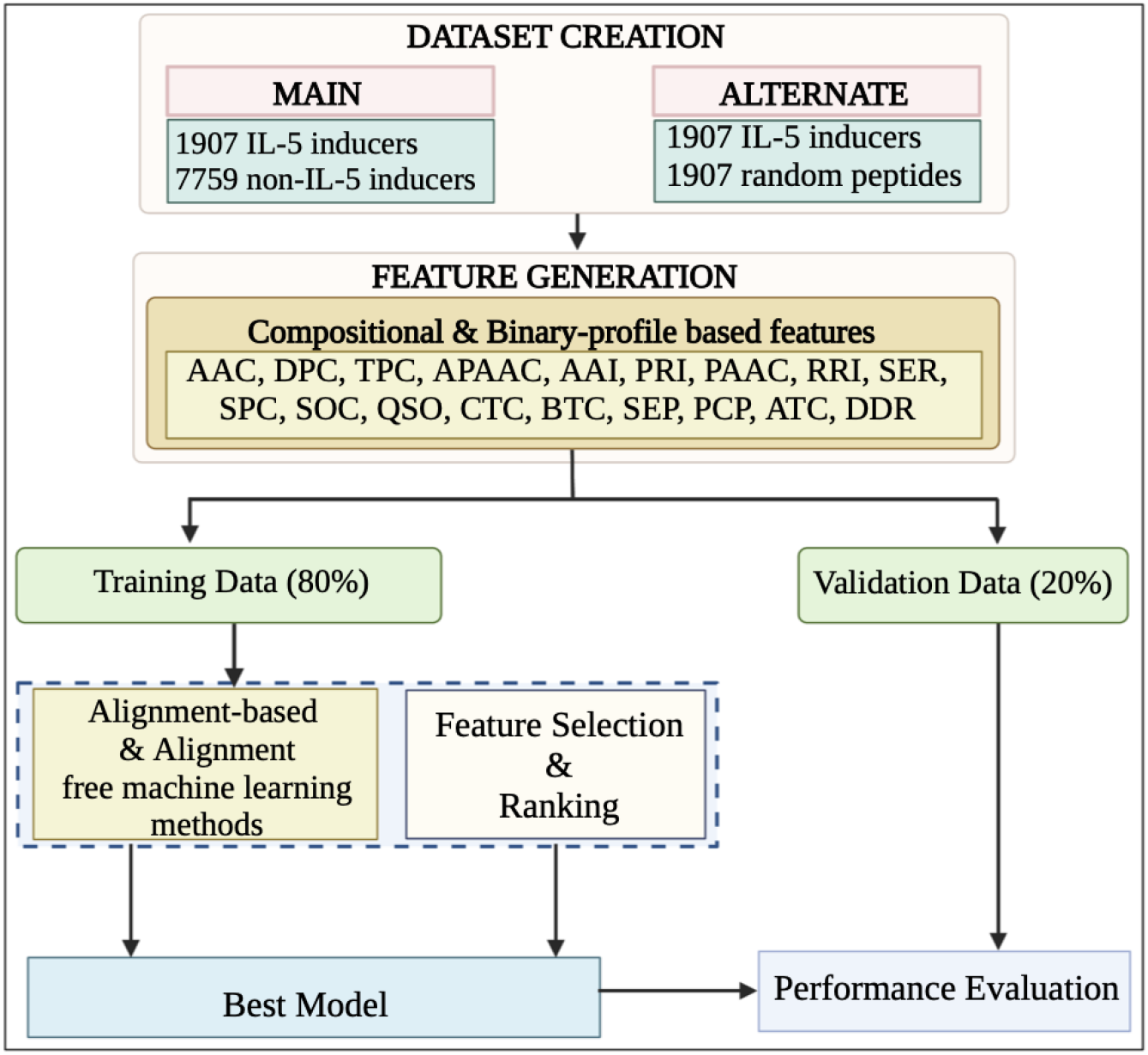
The complete architecture of IL5pred

### Performance evaluation metrics

Model evaluation is a crucial stage in determining the model’s efficiency. Thus, the performance of different ML models was evaluated using standard evaluation metrics, including threshold dependent and independent parameters. The threshold-dependent parameters consist of Accuracy (Acc), Sensitivity (Sens), Specificity (Spec), and Matthews correlation coefficient (MCC), and the Area under the receiver operating characteristic curve (AUC) is a threshold-independent metric. Sens (equation 3) is also known as recall and can be defined as the true positive rate, while Spec (equation 4) is the true negative rate. Acc (equation 5) denotes the percentage of the correctly predicted IL-5 inducers and non-inducers, and MCC (equation 6) is the relation between the predicted and actual values. These performance metrics are commonly used and well-annotated in the previous studies [55,56] and can be calculated as:

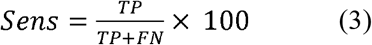

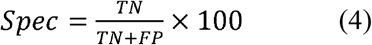

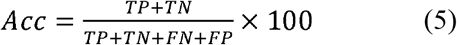

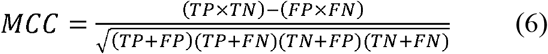

TP, TN, FP, and FN annotate true positive, true negative, false positive, and false negative respectively.

### Hybrid or Ensemble approach

The study explores the potential of a hybrid model that integrates BLAST, motif and alignment-free approach using ML. Initially, BLAST was used to predict the peptide sequence at E-value of 10^−1^. The scores of ‘+0.5’ and‘-0.5’ were assigned for the correct positive and negative predictions respectively, and the score of ‘0’ was provided for no hits. Second, the MERCI program was used to classify the same peptide sequence, and the scores of ‘+0.5’ and ‘-0.5’ were assigned if the motifs were present in positive and negative datasets and 0 for absent. The peptide sequences not identified using BLAST (no hits) and MERCI were further predicted using ML models. Ultimately, the overall score was calculated by combining BLAST, MERCI as well as ML prediction scores. Then, the peptides were assigned as IL-5 inducers and non-IL-5 inducers based on the overall score. This hybrid approach has been applied in different research studies in recent years [54,57].

## Results

### Compositional analysis

The AAC for IL-5 inducing and non-IL-5 inducing peptides was computed. It has been found that the average composition of Phe, Gly, Ile, Lys, Asn, Arg, and Tyr are higher in IL-5 inducing peptides. In contrast, residues like Ala, Thr, Glu, Pro and Val are not preferred in IL-5 inducing peptides. Additionally, we also computed the AAC of random peptides generated from Swiss-Prot. The average AAC for IL-5 inducing, non-IL-5 inducing and random peptides is depicted in Figure 3.

**Figure 3:**
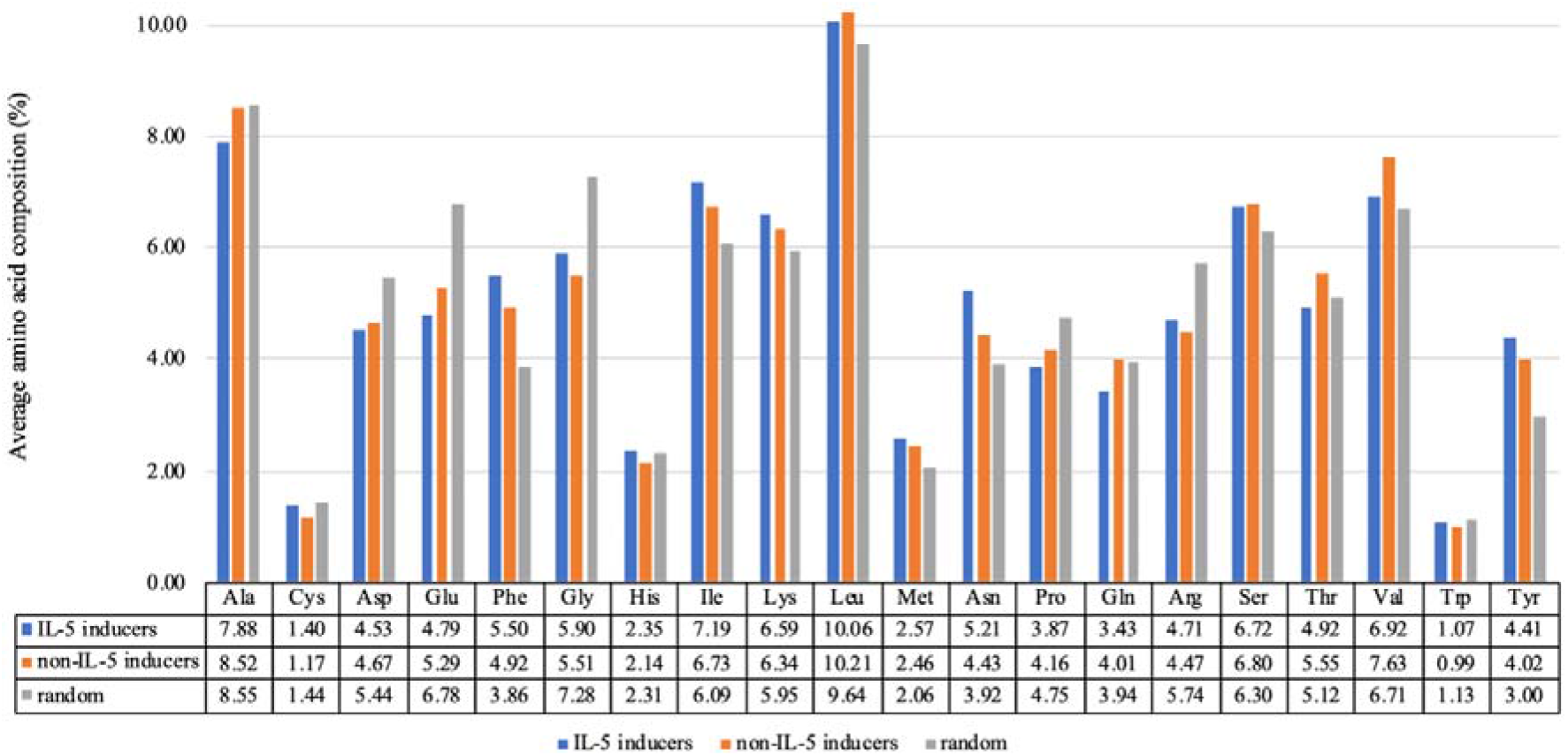
Amino acid composition of IL-5, non-IL-5 and random peptides

### HLA alleles distribution analysis

The distribution of epitopes from IEDB database was also examined for different HLA alleles and is depicted in Figure 4. It has been observed that 9 alleles were found common in both IL-5 inducing and non-IL-5 inducing assays. 24 peptides showed IL-5 inducing whereas only one peptide showed non-IL-5 inducing response against HLA-DRB1*04:01 allele. Similarly, the assays against the HLA-DR allele included 22 IL-5 inducing peptides and 9 non-IL-5 inducing peptides.

**Figure 4:**
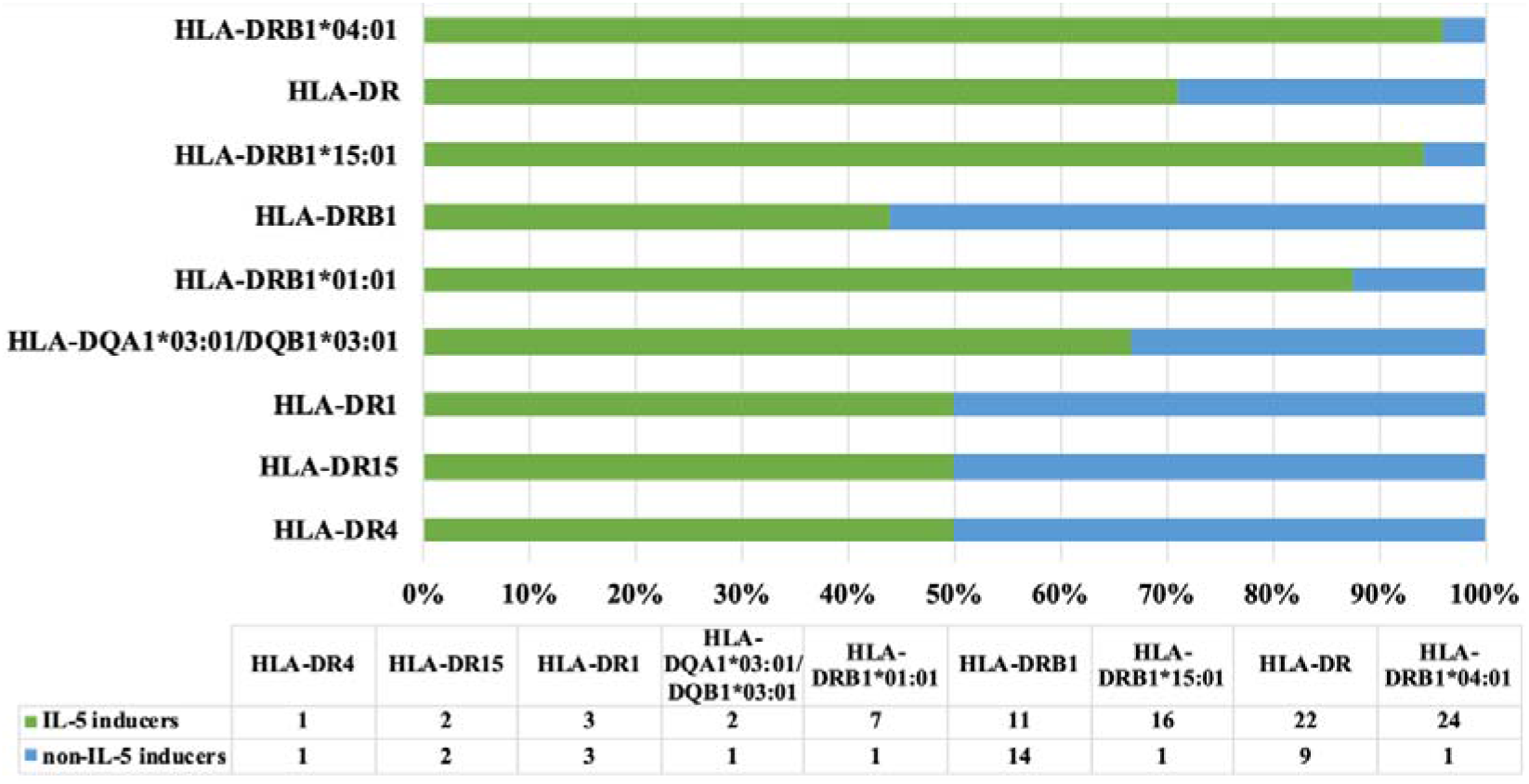
Distribution of HLA alleles among assays reporting IL-5 inducers and IL-5 non-inducers

### Positional preference of residues

We generated the sequence logo for IL-5 inducing peptides to identify the positional preference of individual amino acid residues at specific positions. This analysis exhibited that hydrophobic residues such as Leu, Ile, Val, Phe and Ala are highly predominant in IL-5 inducing peptides. The sequence logo of 9 N-terminal residues of IL-5 inducers is depicted in Figure 5.

**Figure 5:**
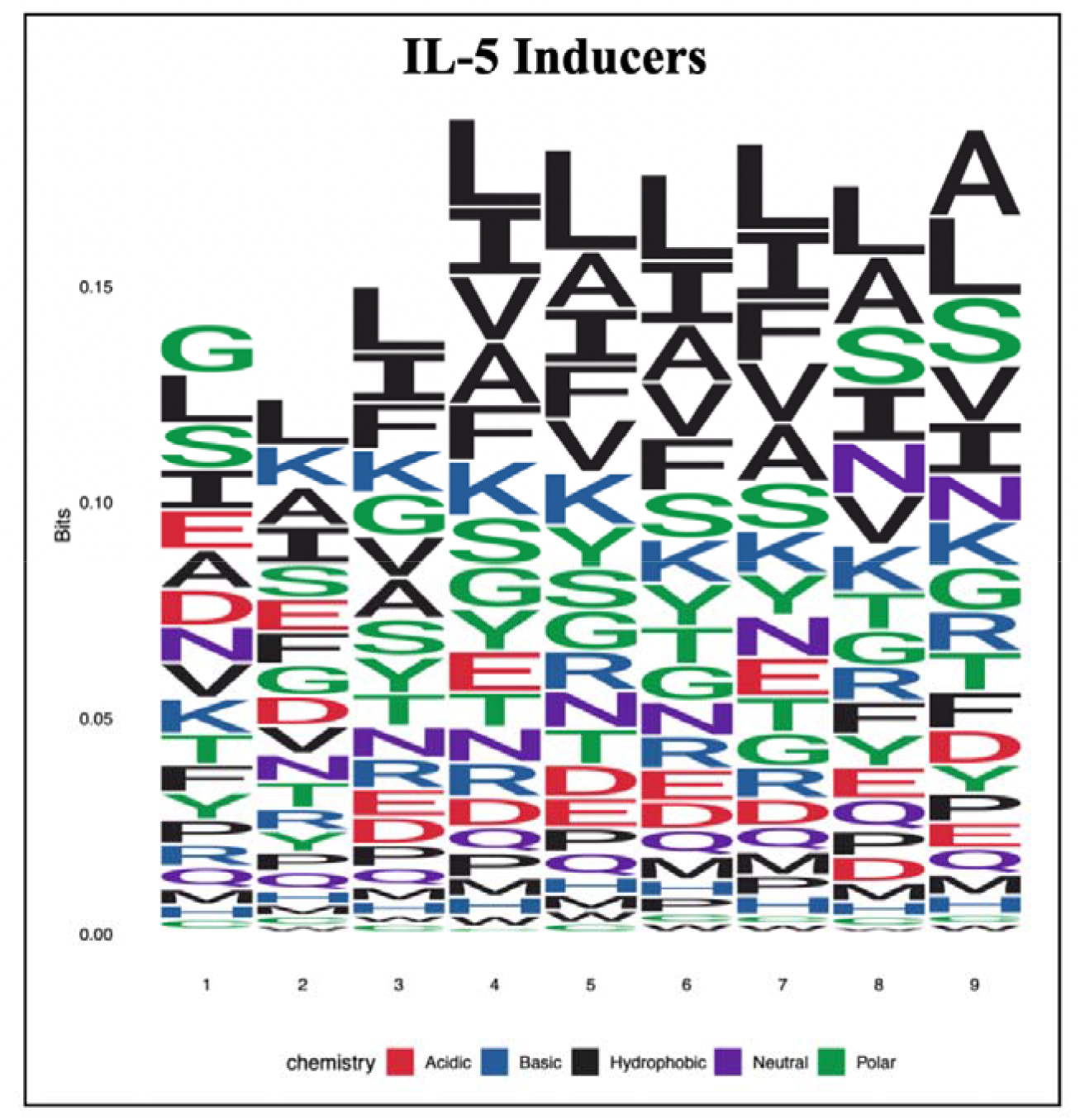
The sequence logo depicting positional conservation of nine amino acid residues at N-terminus for IL-5 inducers

### Alignment-based approach

#### BLAST similarity search

The study utilised BLAST for developing alignment-based models to distinguish between IL-5 inducing and non-IL-5 inducing peptides. A five-fold CV was carried out for evaluating the performance of BLAST. The training dataset was divided into five equal folds, out of which the peptides in four folds were used to build BLAST database, and the sequences in the fifth fold were searched against that database. This process has been iterated five times to avoid any biasness. To evaluate the performance of BLAST on the validation dataset, a BLAST database was generated using all the sequences of the training dataset and then each sequence in the validation dataset was searched against it. The peptide is assigned as IL-5 and non-IL-5 inducer based on the top hit generated by BLAST. For the main dataset, the number of correct hits (sensitivity) rises from 0.13% to 5.56% for the training and 0.1% to 6.72% for the validation datasets. Besides this, it has attained a specificity of 36.39% for training and 40.74% for validation datasets with E-value ranging from 10^−6^ to 10^−1^. As shown in Table 1, the wrong hits (error) increase proportionally with the increase in correct hits for both datasets. Hence, it can be inferred from the result that alignment-based method using BLAST alone is not competent in classifying IL-5 and non-IL-5 inducers since it produces a large number of wrong hits as well as no hits.

**Table 1:** The performance of alignment-based method, developed using BLAST-based similarity on the main dataset

| E-value | Training |  |  |  | Validation |  |  |  |
| --- | --- | --- | --- | --- | --- | --- | --- | --- |
|  | IL-5 inducers |  | non-IL-5 inducers |  | IL-5 inducers |  | non-IL-5 inducers |  |
|  | Chits*<br>(Sens) | Whits<br>(error) | Chits<br>(Spec) | Whits<br>(error) | Chits<br>(Sens) | Whits<br>(error) | Chits<br>(Spec) | Whits<br>(error) |
| $10^{-1}$ | 430<br>(5.56%) | 340<br>(4.4%) | 2814<br>(36.39%) | 354<br>(4.58%) | 130<br>(6.72%) | 99<br>(5.02%) | 788<br>(40.74%) | 97<br>(5.12%) |
| $10^{-2}$ | 249<br>(3.22%) | 207<br>(2.68%) | 1468<br>(18.99%) | 219<br>(2.83%) | 80<br>(4.14%) | 66<br>(3.31%) | 368<br>(19.03%) | 64<br>(3.41%) |
| $10^{-3}$ | 149<br>(1.93%) | 127<br>(1.64%) | 729<br>(9.43%) | 129<br>(1.67%) | 49<br>(2.53%) | 37<br>(1.71%) | 193<br>(9.98%) | 33<br>(1.91%) |
| $10^{-4}$ | 98<br>(1.27%) | 73<br>(0.94%) | 375<br>(4.85%) | 71<br>(0.92%) | 36<br>(1.86%) | 20<br>(0.98%) | 81<br>(4.19%) | 19<br>(1.03%) |
| $10^{-5}$ | 28<br>(0.36%) | 25<br>(0.32%) | 70<br>(0.91%) | 27<br>(0.35%) | 12<br>(0.62%) | 10<br>(0.47%) | 13<br>(0.67%) | 9<br>(0.52%) |
| $10^{-6}$ | 10<br>(0.13%) | 2<br>(0.03%) | 8<br>(0.1%) | 4<br>(0.05%) | 2<br>(0.1%) | 3<br>(0.1%) | 0<br>(0%) | 2<br>(0.16%) |

\*Chits: Correct hits; Whits: Wrong hits; Sens: Sensitivity; Spec: Specificity

### Motif-based approach

In order to extract the motifs solely found in IL-5 inducing and non-IL-5 inducing peptides of the main dataset, we have used the MERCI program. The motifs such as ‘ENSL, LYVGS, HFFN, and NANR’ are exclusively present in IL-5 inducing peptides. Alternatively, ‘VGL, YYA, RSP, and PAG’ motifs are solely found in non-IL-5 inducing peptides. The detailed results are provided in Table 2.

**Table 2:**
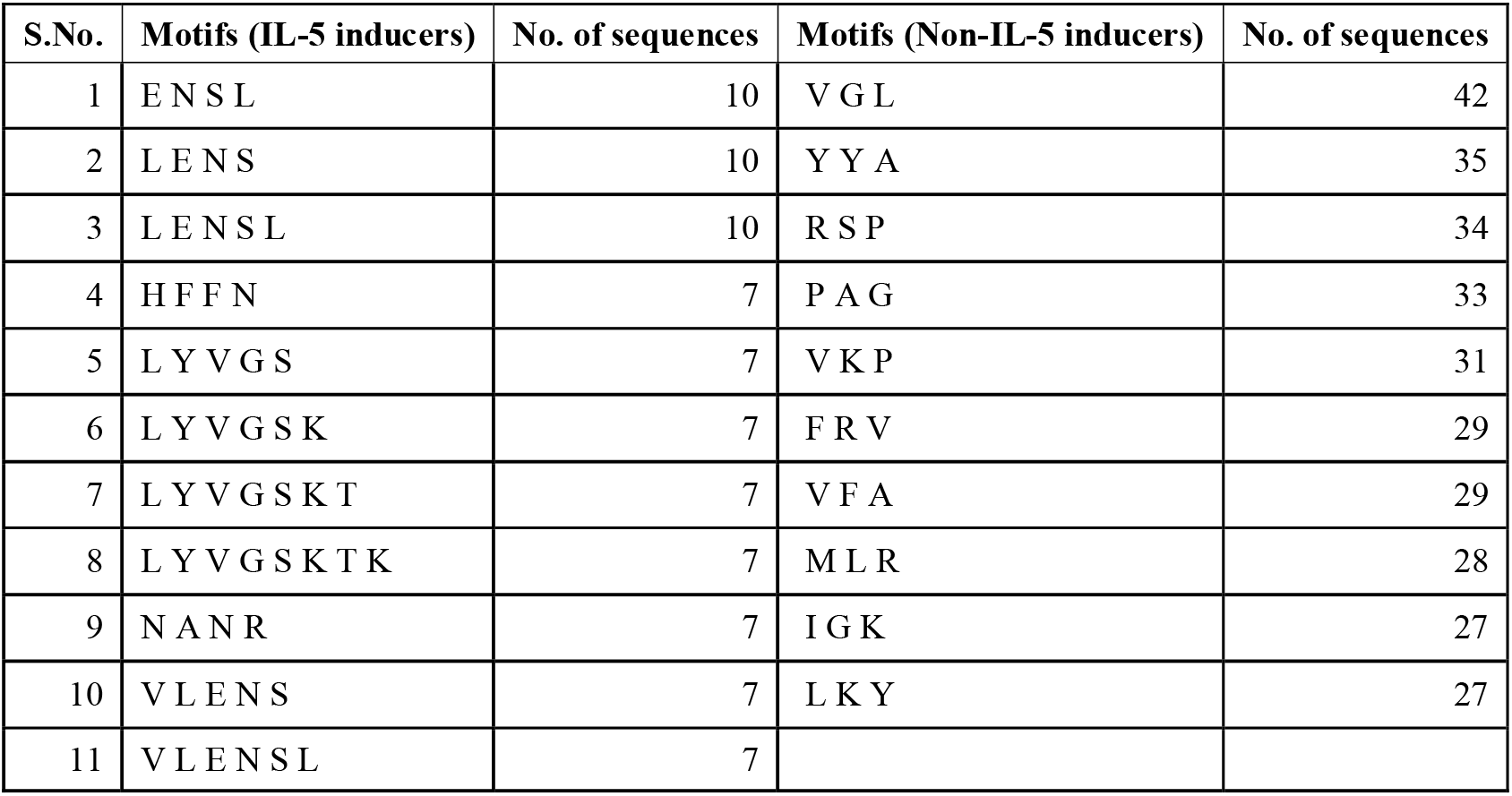
Motifs exclusively present in IL-5 inducing and non-IL-5 inducing peptide sequences

| S.No. | Motifs (IL-5 inducers) | No. of sequences | Motifs (Non-IL-5 inducers) | No. of sequences |
| --- | --- | --- | --- | --- |
| 1 | ENSL | 10 | VGL | 42 |
| 2 | LENS | 10 | YYA | 35 |
| 3 | LENSL | 10 | RSP | 34 |
| 4 | HFFN | 7 | PAG | 33 |
| 5 | LYVGS | 7 | VKP | 31 |
| 6 | LYVGSK | 7 | FRV | 29 |
| 7 | LYVGSKT | 7 | VFA | 29 |
| 8 | LYVGSKTK | 7 | MLR | 28 |
| 9 | NANR | 7 | IGK | 27 |
| 10 | VLENS | 7 | LKY | 27 |
| 11 | VLENSL | 7 |  |  |

### Alignment-free approach

#### Binary-profile based features

We also computed binary-profile-based features to develop alignment-free classification model using several ML techniques. We have built ML models for the N_9_, C_9_ and also for combined terminal residues N_9_C_9_. We found that RF-based model developed using binary profile of C_9_ achieved an AUC of 0.57 on training and 0.58 on validation dataset. RF-model developed using binary profile of N_9_ achieved an AUC of 0.55 on training and 0.57 on validation dataset. In the case of binary profile of N_9_C_9_, XGB-based model achieved an AUC of 0.59 on training and validation datasets. The performance of ML models developed using binary-profile-based features on main dataset is given in Table 3, while the result for the alternate dataset is provided in Table S2. We found that the binary profile-based model does not perform quite well when classifying the peptide sequences.

**Table 3:** The performance of machine learning-based models developed using binary profile of terminal residues of peptides on main dataset

| C<sub>9</sub> |  |  |  |  |  |  |  |  |  |  |
| --- | --- | --- | --- | --- | --- | --- | --- | --- | --- | --- |
| ML | Training |  |  |  |  | Validation |  |  |  |  |
|  | Sens | Spec | Acc | AUC | MCC | Sens | Spec | Acc | AUC | MCC |
| DT | 26.65 | 76.98 | 67.11 | 0.52 | 0.03 | 26.6 | 78.48 | 67.99 | 0.53 | 0.05 |
| RF | 52.9 | 56.19 | 55.55 | 0.57 | 0.07 | 58.82 | 53.27 | 54.4 | 0.58 | 0.1 |
| LR | 51.65 | 53.91 | 53.47 | 0.54 | 0.04 | 54.99 | 52.95 | 53.36 | 0.55 | 0.06 |
| XGB | 53.03 | 54.28 | 54.04 | 0.55 | 0.06 | 55.5 | 52.17 | 52.84 | 0.55 | 0.06 |
| KNN | 67.15 | 36.65 | 42.63 | 0.53 | 0.03 | 67.01 | 36.49 | 42.66 | 0.53 | 0.03 |
| GNB | 53.03 | 52.09 | 52.28 | 0.53 | 0.04 | 58.31 | 50.49 | 52.07 | 0.57 | 0.07 |
| **N<sub>9</sub>** |  |  |  |  |  |  |  |  |  |  |
| DT | 50.59 | 51.43 | 51.27 | 0.52 | 0.02 | 58.31 | 44.98 | 47.67 | 0.53 | 0.03 |
| RF | 50.53 | 55.84 | 54.8 | 0.55 | 0.05 | 54.73 | 53.79 | 53.98 | 0.57 | 0.07 |
| LR | 47.89 | 57.08 | 55.28 | 0.54 | 0.04 | 44.5 | 55.8 | 53.52 | 0.52 | 0 |
| XGB | 52.77 | 53.27 | 53.17 | 0.54 | 0.05 | 51.15 | 51.72 | 51.6 | 0.52 | 0.02 |
| KNN | 49.87 | 56.66 | 55.33 | 0.55 | 0.05 | 53.2 | 53.66 | 53.57 | 0.57 | 0.06 |
| GNB | 50.86 | 51.05 | 51.01 | 0.52 | 0.02 | 54.22 | 46.66 | 48.19 | 0.52 | 0.01 |
| **N<sub>9</sub>C<sub>9</sub>** |  |  |  |  |  |  |  |  |  |  |
| DT | 24.27 | 78.88 | 68.17 | 0.52 | 0.03 | 27.88 | 78.61 | 68.36 | 0.53 | 0.06 |
| RF | 57.65 | 54.1 | 54.8 | 0.59 | 0.09 | 62.4 | 50.23 | 52.69 | 0.58 | 0.1 |
| LR | 53.5 | 52.61 | 52.78 | 0.54 | 0.05 | 59.08 | 52.11 | 53.52 | 0.56 | 0.09 |
| XGB | 54.95 | 56.93 | 56.54 | 0.59 | 0.1 | 55.75 | 55.02 | 55.17 | 0.59 | 0.09 |
| KNN | 66.56 | 40.44 | 45.56 | 0.56 | 0.06 | 62.4 | 40.31 | 44.78 | 0.54 | 0.02 |
| GNB | 51.65 | 52.56 | 52.38 | 0.53 | 0.03 | 59.08 | 48.74 | 50.83 | 0.56 | 0.06 |

\*Sens: Sensitivity; Spec: Specificity; Acc: Accuracy; MCC: Matthews correlation coefficient; AUC: Area under receiver operating characteristic curve; RF: Random Forest; LR: Logistic Regression; XGB: eXtreme Gradient Boosting; KNN: K-nearest neighbour; GNB: Gaussian Naïve Bayes; DT: Decision Tree

### All compositional features

A vector of 9553 features was computed against each sequence for both main and alternate datasets using Pfeature standalone tool [42]. We computed composition-based features including AAC, DPC, TPC and others to distinguish IL-5 and non-IL-5 inducing peptides. These 9553 features were used to develop alignment-free classification model using several ML techniques. The performance of ML models developed using all compositional features on the main dataset is listed in Table 4. We found that LR-based model attained an AUC of 0.68 on training and validation datasets. In addition, the performance of ML models developed using all compositional features on the alternate dataset is tabulated in Tables S3.

**Table 4:** The performance of machine learning models developed using all compositional features on the main dataset

|  | Training |  |  |  |  | Validation |  |  |  |  |
| --- | --- | --- | --- | --- | --- | --- | --- | --- | --- | --- |
| ML | Sens | Spec | Acc | AUC | MCC | Sens | Spec | Acc | AUC | MCC |
| DT | 52.97 | 53.02 | 53.01 | 0.55 | 0.05 | 80.31 | 29.29 | 39.61 | 0.57 | 0.09 |
| RF | 61.74 | 62.61 | 62.44 | 0.68 | 0.20 | 61.38 | 59.17 | 59.62 | 0.67 | 0.17 |
| LR | 63.65 | 64.24 | 64.12 | 0.68 | 0.23 | 64.19 | 62.22 | 62.62 | 0.68 | 0.21 |
| XGB | 62.14 | 59.25 | 59.82 | 0.65 | 0.17 | 62.66 | 58.78 | 59.57 | 0.65 | 0.17 |
| KNN | 53.63 | 54.09 | 54.00 | 0.56 | 0.06 | 61.13 | 50.94 | 53.00 | 0.58 | 0.10 |
| GNB | 41.29 | 75.87 | 69.09 | 0.59 | 0.15 | 48.08 | 76.28 | 70.58 | 0.62 | 0.22 |

\*Sens: Sensitivity; Spec: Specificity; Acc: Accuracy; MCC: Matthews correlation coefficient; AUC: Area under receiver operating characteristic curve; RF: Random Forest; LR: Logistic Regression; XGB: eXtreme Gradient Boosting; KNN: K-nearest neighbour; GNB: Gaussian Naïve Bayes; DT: Decision Tree

Alternatively, SVC-L1 method was then used to reduce these features to 318 (main dataset) and 164 (alternate dataset). These selected features for the main dataset were utilised to build several prediction models and their performance is tabulated in Table 5. The performance of machine learning models developed using selected features on the alternate dataset is shown in Table S4.

**Table 5:** The performance of machine learning models developed using selected features on the main dataset

| 318 Features |  |  |  |  |  |  |  |  |  |  |  |
| --- | --- | --- | --- | --- | --- | --- | --- | --- | --- | --- | --- |
| ML | Training |  |  |  |  |  | Validation |  |  |  |  |
|  | Threshold | Sens | Spec | Acc | AUC | MCC | Sens | Spec | Acc | AUC | MCC |
| DT | 0.48 | 57.98 | 53.96 | 54.75 | 0.58 | 0.10 | 49.87 | 60.60 | 58.43 | 0.56 | 0.09 |
| RF | 0.19 | 69.86 | 65.03 | 65.97 | 0.74 | 0.28 | 68.80 | 59.88 | 61.69 | 0.70 | 0.23 |
| LR | 0.48 | 67.41 | 66.92 | 67.02 | 0.74 | 0.28 | 61.64 | 62.87 | 62.62 | 0.67 | 0.20 |
| XGB | 0.17 | 63.65 | 63.05 | 63.17 | 0.70 | 0.22 | 62.40 | 61.05 | 61.32 | 0.67 | 0.19 |
| KNN | 0.18 | 63.52 | 59.20 | 60.05 | 0.65 | 0.18 | 61.89 | 56.32 | 57.45 | 0.62 | 0.15 |
| GNB | 1 | 93.27 | 29.46 | 41.97 | 0.61 | 0.21 | 85.68 | 27.48 | 39.25 | 0.57 | 0.12 |

\*Sens: Sensitivity; Spec: Specificity; Acc: Accuracy; MCC: Matthews correlation coefficient; AUC: Area under receiver operating characteristic curve; RF: Random Forest; LR: Logistic Regression; XGB: eXtreme Gradient Boosting; KNN: K-nearest neighbour; GNB: Gaussian Naïve Bayes; DT: Decision Tree

Further, we ranked these reduced features using feature-selector tool based on their normalized and cumulative importance. The detailed results for the top-ranked features are tabulated in Table S5. Based on these top-ranked features (10, 50, 100, 150,….), several classification models were developed. The performance of the best models developed using the top-ranked features on the main dataset is listed in Table 6. In addition, the performance of other classification models developed using these ranked features on the main and alternate datasets was also computed and tabulated in Tables S6 & S7.

**Table 6:** Performance of the best random forest-based models developed on top-ranked features of the main dataset

|  | Training |  |  |  |  | Validation |  |  |  |  |
| --- | --- | --- | --- | --- | --- | --- | --- | --- | --- | --- |
| Features | Sens | Spec | Acc | AUC | MCC | Sens | Spec | Acc | AUC | MCC |
| Top10 | 54.02 | 56.90 | 56.34 | 0.59 | 0.09 | 57.55 | 55.54 | 55.95 | 0.60 | 0.11 |
| Top50 | 64.18 | 59.44 | 60.37 | 0.67 | 0.19 | 64.19 | 56.12 | 57.76 | 0.65 | 0.16 |
| Top100 | 62.53 | 65.15 | 64.64 | 0.69 | 0.22 | 62.66 | 63.38 | 63.24 | 0.67 | 0.21 |
| Top150 | 64.51 | 64.11 | 64.19 | 0.71 | 0.23 | 66.24 | 60.73 | 61.84 | 0.70 | 0.22 |
| Top200 | 64.91 | 66.06 | 65.83 | 0.73 | 0.25 | 65.47 | 62.67 | 63.24 | 0.70 | 0.23 |

\*Sens: Sensitivity; Spec: Specificity; Acc: Accuracy; MCC: Matthews correlation coefficient; AUC: Area under receiver operating characteristic curve; RF: Random Forest; LR: Logistic Regression; XGB: eXtreme Gradient Boosting; KNN: K-nearest neighbour; GNB: Gaussian Naïve Bayes; DT: Decision Tree

### Different types of compositional features

These compositional features were used to develop alignment-free classification model using several ML techniques. The performance of the best models developed using different composition-based features on the main dataset is listed in Table 7. Among all, we found that DPC-based RF model achieved an AUC of 0.74 on training and validation datasets. It could be inferred from the results that alignment-free ML-based model using DPC has performed significantly better than other compositional-based features models. Furthermore, the performance of other classification models developed using compositional-based features on the main and alternate datasets was computed and tabulated in Supplementary Tables S8 & S9.

**Table 7:** Performance of best machine learning-based models developed on compositional-based features of the main dataset

| Features | ML | Training |  |  |  |  | Validation |  |  |  |  |
| --- | --- | --- | --- | --- | --- | --- | --- | --- | --- | --- | --- |
|  |  | Sens | Spec | Acc | AUC | MCC | Sens | Spec | Acc | AUC | MCC |
| AAC | RF | 61.61 | 61.73 | 61.71 | 0.67 | 0.19 | 63.17 | 59.69 | 60.39 | 0.66 | 0.19 |
| AAI | RF | 59.57 | 58.72 | 58.89 | 0.63 | 0.15 | 60.61 | 56.38 | 57.24 | 0.62 | 0.14 |
| APAAC | RF | 63.26 | 61.68 | 61.99 | 0.68 | 0.20 | 60.10 | 59.37 | 59.51 | 0.65 | 0.16 |
| ATC | GNB | 54.29 | 51.98 | 52.43 | 0.54 | 0.05 | 57.55 | 47.89 | 49.85 | 0.54 | 0.04 |
| BTC | XGB | 49.54 | 60.28 | 58.17 | 0.57 | 0.08 | 49.11 | 59.88 | 57.70 | 0.56 | 0.07 |
| CTC | RF | 59.76 | 59.57 | 59.61 | 0.64 | 0.16 | 62.40 | 56.77 | 57.91 | 0.65 | 0.15 |
| DDR | RF | 56.86 | 59.27 | 58.80 | 0.62 | 0.13 | 59.59 | 56.90 | 57.45 | 0.62 | 0.13 |
| **DPC** | **RF** | **68.01** | **66.20** | **66.56** | **0.74** | **0.28** | **68.80** | **66.11** | **66.65** | **0.74** | **0.29** |
| PAAC | RF | 61.21 | 60.65 | 60.76 | 0.67 | 0.18 | 63.17 | 60.53 | 61.07 | 0.66 | 0.19 |
| PCP | RF | 58.58 | 62.56 | 61.78 | 0.66 | 0.17 | 58.31 | 61.24 | 60.65 | 0.66 | 0.16 |
| PRI | RF | 55.34 | 58.67 | 58.02 | 0.60 | 0.11 | 52.43 | 55.74 | 55.07 | 0.56 | 0.07 |
| QSO | RF | 62.01 | 57.26 | 58.19 | 0.64 | 0.15 | 63.17 | 54.18 | 56.00 | 0.62 | 0.14 |
| RRI | RF | 56.99 | 60.10 | 59.49 | 0.62 | 0.14 | 54.73 | 57.29 | 56.77 | 0.59 | 0.10 |
| SEP | DT | 51.19 | 58.41 | 57.00 | 0.57 | 0.08 | 53.45 | 52.60 | 52.74 | 0.56 | 0.05 |
| SER | RF | 61.94 | 60.76 | 60.99 | 0.67 | 0.18 | 60.87 | 60.34 | 60.45 | 0.66 | 0.17 |
| SOC | LR | 53.83 | 49.36 | 50.23 | 0.52 | 0.03 | 52.69 | 48.74 | 49.54 | 0.51 | 0.01 |
| SPC | RF | 61.28 | 58.70 | 59.21 | 0.65 | 0.16 | 60.10 | 60.01 | 60.03 | 0.66 | 0.16 |
| TPC | RF | 68.67 | 59.86 | 61.59 | 0.71 | 0.23 | 71.10 | 58.59 | 61.12 | 0.71 | 0.24 |

\*Sens: Sensitivity; Spec: Specificity; Acc: Accuracy; MCC: Matthews correlation coefficient; AUC: Area under receiver operating characteristic curve; RF: Random Forest; LR: Logistic Regression; XGB: eXtreme Gradient Boosting; KNN: K-nearest neighbour; GNB: Gaussian Naïve Bayes; DT: Decision Tree

### Cascade machine-learning model

In addition, we employed a bilayer cascade ML-based approach to distinguish IL-5 and non-IL-5 with improved accuracy. In this approach, the prediction was carried out using two layers of machine learning algorithms. In the first layer, 108 ML models were generated by amalgamating the prediction scores of 18 different types of features computed using six different ML techniques. The second layer has been trained on the scores generated by the first layer. Hence, the second layer learns from the results of the first layer classifiers and produces a final cascade ML model [58] (Figure 6). We found that among all the ML models, XGB-based classifier attained an AUC of 0.69 on training and 0.67 on validation dataset for the main dataset (Table S10). The performance of the alternate dataset using cascade approach is given in Table S10.

**Figure 6:**
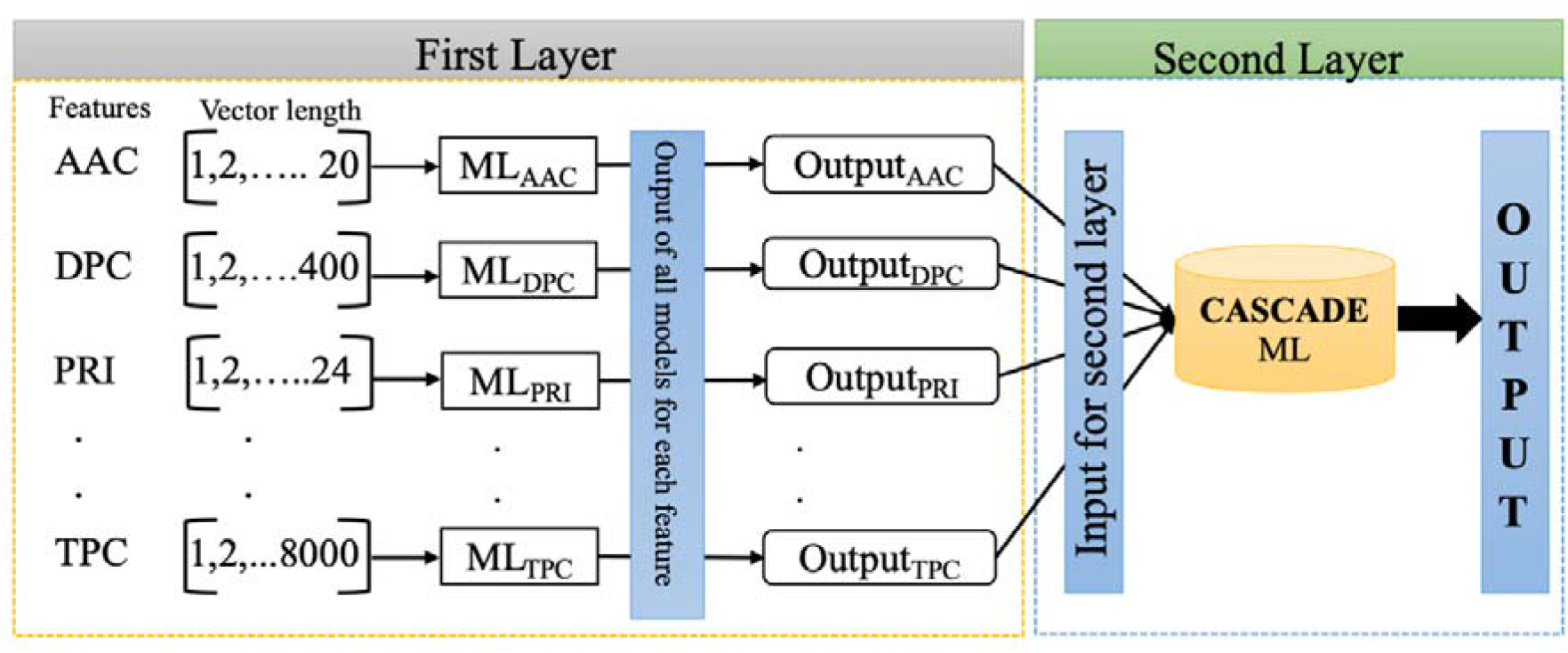
The diagrammatic representation of bilayer cascade ML-based approach

### Selected dipeptide compositional features

Since we achieved the best performance on DPC features using RF-based model, thus we utilised these features to enhance the classification performance. Feature-selector tool was used to rank these features based on their importance. Using these top-ranked features, we developed several machine learning-based models and their performances are tabulated in Table 8. We observed that RF-based model on DPC-250 features attains the best performance with an AUC of 0.74 on training and 0.75 on validation dataset respectively, with balanced sensitivity and specificity for main dataset. The same procedure was carried out on alternate dataset and the detailed results are provided in Supplementary Tables S11.

**Table 8:** The performance of best machine learning-based models developed using composition of selected dipeptides

| Features | Training |  |  |  |  | Validation |  |  |  |  |
| --- | --- | --- | --- | --- | --- | --- | --- | --- | --- | --- |
|  | Sens | Spec | Accuracy | AUC | MCC | Sens | Spec | Accuracy | AUC | MCC |
| Top50 | 58.64 | 60.15 | 59.86 | 0.63 | 0.15 | 59.34 | 60.86 | 60.55 | 0.64 | 0.16 |
| Top100 | 63.33 | 63.11 | 63.15 | 0.69 | 0.21 | 67.52 | 63.19 | 64.06 | 0.72 | 0.25 |
| Top150 | 64.05 | 66.89 | 66.34 | 0.72 | 0.25 | 69.57 | 65.91 | 66.65 | 0.74 | 0.29 |
| Top200 | 64.58 | 68.42 | 67.67 | 0.73 | 0.27 | 65.22 | 70.06 | 69.08 | 0.74 | 0.29 |
| **Top250** | **65.70** | **68.21** | **67.72** | **0.74** | **0.28** | **66.24** | **69.15** | **68.56** | **0.75** | **0.29** |
| Top300 | 68.34 | 65.01 | 65.66 | 0.74 | 0.27 | 71.36 | 65.65 | 66.81 | 0.75 | 0.30 |
| Top350 | 67.81 | 66.72 | 66.93 | 0.74 | 0.28 | 68.29 | 67.14 | 67.37 | 0.75 | 0.29 |
| All400 | 68.01 | 66.20 | 66.56 | 0.74 | 0.28 | 68.80 | 66.11 | 66.65 | 0.74 | 0.29 |

\*Sens: Sensitivity; Spec: Specificity; Acc: Accuracy; MCC: Matthews correlation coefficient; AUC: Area under receiver operating characteristic curve; RF: Random Forest; LR: Logistic Regression; XGB: eXtreme Gradient Boosting; KNN: K-nearest neighbour; GNB: Gaussian Naïve Bayes; DT: Decision Tree

We developed several ML-based methods to predict IL-5 using compositional and binary profile-based features computed from the peptide sequence. We could infer from the results that alignment-free ML-based model using DPC-250 features has performed significantly better than other feature-based models. Hence, in the current study, the hybrid model was developed using DPC-250 features of the peptides.

### Hybrid approach

To combat the shortcomings of individual techniques, we have explored the potential of our hybrid method that combines BLAST, motif with ML using DPC-250 features. This approach was built to classify the IL-5 inducers with high precision. For this, the composition-based model is integrated with MERCI and BLAST-based approaches.

Followed by the MERCI approach, peptides were distinguished using BLAST with an E-value of 10^−1^. The sequence is assigned as IL-5 and non-IL-5 inducers based on the top hit of BLAST, while the peptides that are unpredicted by BLAST were further predicted using DPC-based ML model. This hybrid method considerably improved the accuracy on validation dataset, hence overcoming the limitation of each method as shown in Table 9. RF-based model performed the best on main dataset and achieved AUC of 0.92 and 0.94 on the training and validation dataset. The results for the alternate dataset are provided in Supplementary Table S12.

**Table 9:** Performance of hybrid approach that combines BLAST, motif and selected dipeptide composition-based models on the main dataset

| ML | Threshold | Training |  |  |  |  | Validation |  |  |  |  |
| --- | --- | --- | --- | --- | --- | --- | --- | --- | --- | --- | --- |
|  |  | Sens | Spec | Acc | AUC | MCC | Sens | Spec | Acc | AUC | MCC |
| RF | 0.2 | 79.68 | 82.14 | 81.66 | 0.92 | 0.54 | 82.86 | 84.71 | 84.33 | 0.94 | 0.60 |
| DT | 0.38 | 76.52 | 76.66 | 76.63 | 0.83 | 0.45 | 82.35 | 78.55 | 79.32 | 0.86 | 0.52 |
| GNB | 0.5 | 79.29 | 74.97 | 75.81 | 0.82 | 0.45 | 81.59 | 78.22 | 78.90 | 0.84 | 0.51 |
| KNN | 0.24 | 78.63 | 80.04 | 79.76 | 0.91 | 0.50 | 76.47 | 84.06 | 82.52 | 0.92 | 0.54 |
| XGB | 0.16 | 79.55 | 79.63 | 79.62 | 0.91 | 0.50 | 83.12 | 81.66 | 81.95 | 0.93 | 0.56 |
| LR | 0.47 | 78.3 | 79.50 | 79.27 | 0.89 | 0.49 | 81.07 | 80.43 | 80.56 | 0.92 | 0.53 |

\*Sens: Sensitivity; Spec: Specificity; Acc: Accuracy; MCC: Matthews correlation coefficient; AUC: Area under receiver operating characteristic curve; RF: Random Forest; LR: Logistic Regression; XGB: eXtreme Gradient Boosting; KNN: K-nearest neighbour; GNB: Gaussian Naïve Bayes; DT: Decision Tree

### Implementation of IL5pred server

A user-friendly web server IL5pred was developed for predicting IL-5 inducing peptides with better efficiency. The main components of the web server are Predict, Design, Protein scan, Motif scan, and Blast scan. The modules are explained below.

i. **Predict Module**: Enables the user to submit single/multiple peptide sequences in FASTA format and will predict the peptides as IL-5 inducers or non-IL-5 inducers. The module will provide the user prediction scores and results (IL-5 inducers/ non-inducers) based on the chosen threshold value.
ii. **Design Module**: Enables the user to design novel IL-5 inducers with better activity. The user can provide the input sequence in single-line format. It will generate all possible mutants peptides with a single mutation which are further predicted using the model.
iii. **Protein scan**: Allows the user to identify IL-5 inducing regions in the given protein sequence by generating overlapping patterns based on the chosen length.
iv. **Motif scan**: Allows the user to search for the motifs present in IL-5 inducing peptide sequences. This module used MERCI program to extract motifs from the given sequence.
v. **Blast scan**: Uses an alignment-based search method, BLAST. The module allows to search the query sequence against the database of known IL-5 inducing peptides and assign it as IL-5 inducers based on the match from the database.

The web server can be accessed at (https://webs.iiitd.edu.in/raghava/il5pred/). The python-based standalone package of IL5pred was also developed which can be downloaded from (https://webs.iiitd.edu.in/raghava/il5pred/stand.php). The web server is user-friendly and compatible with all contemporary gadgets, such as tablets, and desktops.

## Discussion

IL-5 is an eosinophil regulatory cytokine involved in the maturation, differentiation, activation and migration of eosinophils [5,59]. It plays a role in a wide variety of diseases such as asthma, dermatitis, eosinophilic esophagitis, and hyper-eosinophilic syndrome [12]. Several autoimmune disorders, such as Hashimoto’s thyroiditis and Graves’ disease, are also associated with elevated IL-5 levels and eosinophilic infiltration [60]. A recent study by Lucas et al. reported that the level of IL-5 is found to be elevated in patients with severe COVID-19 conditions [61]. Past studies have revealed the role of IL-5 in treating several eosinophil-mediated disorders [62] and possess anti-tumor activity. For instance, Ikutani et al. have shown the role of IL-5 inducing cells to control eosinophil infiltration in lungs hence preventing tumor metastasis [63]. IL-5 can act as an appealing therapeutic target due to its pivotal role in the majority of eosinophil-mediated diseases [64–66]. Currently, three FDA-approved anti-IL-5 therapeutic agents, such as benralizumab, mepolizumab and reslizumab, are available in the market [9].

Taking into consideration, the role of IL-5 in several diseases there is a necessity to develop a computational method that can predict IL-5 inducing peptides. In the current study, a prediction method, named IL5pred that can be used to anticipate the peptides as IL-5 inducers and non-IL-5 inducers. For this, we have created two datasets namely main and alternate. The main dataset comprises experimentally validated 1907 IL-5 inducing and 7759 non-IL-5 inducing peptides obtained from IEDB. While the alternate dataset contains 1907 IL-5 inducing peptides from main dataset and 1907 random peptides generated from Swiss-Prot. The compositional analysis shows that IL-5 inducing peptides are abundant in Phe, Gly, Ile, Lys, Asn, Arg, and Tyr. Besides composition, the order of the residue is an essential feature and plays a vital role in defining its activity. For this, we also analyzed the residue preference and observed that hydrophobic residues such as Leu, Ile, Val, Phe and Ala are highly preferred at N-terminus of IL-5 inducing peptides. Along with this, we have extracted the motifs exclusively present in IL-5 inducing peptides. Moreover, a number of sequence-based features including compositional and binary profiles were calculated for the peptides using Pfeature standalone tool. We have computed a total of 9553 features and further reduced them by SVC-L1 feature selection method and ranked using the feature selector tool. Utilizing these features, we have developed numerous prediction models using alignment-free ML techniques. We have also employed a bilayer cascade ML-based approach to distinguish IL-5 and non-IL-5 with improved accuracy. We tried every possible feature in an effort to accurately predict IL-5. However, ML-based models that used DPC-250 features outperformed other feature-based models.

Apart from this, commonly used alignment-based method, BLAST was also employed to annotate the peptide sequence. If the query sequence shows a high degree of similarity with a known peptide, then the same function is assigned to the query peptide. It is found from Table 1 that BLAST can predict IL-5 inducers, but it also produces an enormous number of no hits. To cope with this limitation, alignment-based, and alignment-free approaches using DPC-250 were combined to develop an improved method. We observed that the hybrid model outperformed other models in the case of main dataset. Nonetheless, the hybrid model based on DPC-250, motif and BLAST can accurately distinguish IL-5 and non-IL-5 inducing peptides in terms of accuracy. To expedite the research community working in the field of immunotherapy, we have incorporated the best-performing model into the web server. IL5pred is a user-friendly, freely accessible web server which is compatible with laptops and desktops. We expect that researchers will use our prediction method to develop more accurate peptide-based therapeutics for a variety of diseases.

### Limitation of the study

This is the first systematic attempt to develop *in silico* models for identifying IL-5 inducing peptides. A large enough dataset is required to develop a more robust and accurate classification method. One limitation of this method is the limited number of experimentally validated IL-5 inducing peptides. We have made an effort to create more accurate and reliable method while keeping this constraint in mind.

## Author Contributions

NLD collected, compiled, and processed the data sets. NLD and NS developed computer programs. NLD and NS implemented the algorithms and prediction models. NLD and NS created the web server. NLD and NS analysed the results. NLD, NS and GPSR wrote the manuscript. GPSR conceived and coordinated the project and provided overall supervision of the project. All authors have read and approved the final manuscript.

## Acknowledgement

We are thankful to the Department of Biotechnology (DBT) for providing an infrastructure grant to the institute. NLD is thankful to DBT-RA program in Biotechnology and Life Sciences for providing Research Associate fellowship. NS is thankful to the Department of Science and Technology (DST-INSPIRE) for providing Senior Research Fellowship. NLD, and NS are thankful to Department of Computational Biology, IIIT-Delhi for infrastructure and facilities. We would like to acknowledge that Figures were created using BioRender.com.

## Data Availability Statement

All the datasets generated for this study are available at the “IL5pred” webserver, https://webs.iiitd.edu.in/raghava/il5pred/stand.php. The source code is hosted on GitHub and can be found at https://github.com/raghavagps/il5pred.

